# Neural Computations Underlying Causal Structure Learning

**DOI:** 10.1101/228593

**Authors:** Momchil S. Tomov, Hayley M. Dorfman, Samuel J. Gershman

## Abstract

Behavioral evidence suggests that beliefs about causal structure constrain associative learning, determining which stimuli can enter into association, as well as the functional form of that association. Bayesian learning theory provides one mechanism by which structural beliefs can be acquired from experience, but the neural basis of this mechanism is unknown. A recent study (Gershman, 2017) proposed a unified account of the elusive role of “context” in animal learning based on Bayesian updating of beliefs about the structure of causal relationships between contexts and cues in the environment. The model predicts that the computations which arbitrate between these abstract causal structures are distinct from the computations which learn the associations between particular stimuli under a given structure. In this study, we used fMRI with male and female human subjects to interrogate the neural correlates of these two distinct forms of learning. We show that structure learning signals are encoded in rostrolateral prefrontal cortex and the angular gyrus, anatomically distinct from correlates of associative learning. Within-subject variability in the encoding of these learning signals predicted variability in behavioral performance. Moreover, representational similarity analysis suggests that some regions involved in both forms of learning, such as parts of the inferior frontal gyrus, may also encode the full probability distribution over causal structures. These results provide evidence for a neural architecture in which structure learning guides the formation of associations.

**Significance Statement:** Animals are able to infer the hidden structure behind causal relations between stimuli in the environment, allowing them to generalize this knowledge to stimuli they have never experienced before. A recently published computational model based on this idea provided a parsimonious account of a wide range of phenomena reported in the animal learning literature, suggesting that the neural mechanisms dedicated to learning this hidden structure are distinct from those dedicated to acquiring particular associations between stimuli. Here we validate this model by measuring brain activity during a task which dissociates structure learning from associative learning. We show that different brain networks underlie the two forms of learning and that the neural signal corresponding to structure learning predicts future behavioral performance.

## Introduction

Classical learning theories posit that animals learn associations between sensory stimuli and rewarding outcomes (Rescorla and Wagner, 1972; Pearce and Bouton, 2001). These theories have achieved remarkable success in explaining a wide range of behaviors using simple mathematical rules. Yet numerous studies have challenged some of their foundational premises (Miller et al., 1995; Gershman et al., 2015; Dunsmoor et al., 2015). One particularly longstanding puzzle for these theories is the multifaceted role of contextual stimuli in associative learning. Some studies have shown that the context in which learning takes place is largely irrelevant (Bouton and King, 1983; Lovibond et al., 1984; Kaye et al., 1987; Bouton and Peck, 1989), whereas others have found that context plays the role of an “occasion setter,” modulating cue-outcome associations without itself acquiring associative strength (Swartzentruber and Bouton, 1986; Grahame et al., 1990; Bouton and Bolles, 1993; Swartzentruber, 1995). Yet other studies suggest that context acts like another punctate cue, entering into summation and cue competition with other stimuli (Balaz et al., 1981; Grau and Rescorla, 1984). The multiplicity of such behavioral patterns defies explanation in terms of a single associative structure, suggesting instead that different structures may come into play depending on the task and training history.

Computational modeling has begun to unravel this puzzle, using the idea that structure is a latent variable inferred from experience (Gershman, 2017). On this account, each structure corresponds to a causal model of the environment, specifying the links between context, cues and outcomes, as well as their functional form (modulatory vs. additive). The learner thus faces the joint problem of inferring both the structure and the strength of causal relationships, which can be implemented computationally using Bayesian learning (Griffiths and Tenenbaum, 2005; Körding et al., 2007; Meder et al., 2014). This account can explain why different tasks and training histories produce different forms of context-dependence: variations across tasks induce different probabilistic beliefs about causal structure. For example, Gershman (2017) showed that manipulations of context informativeness, outcome intensity, and number of training trials have predictable effects on the functional role of context in animal learning experiments (Odling-Smee, 1978; Preston et al., 1986).

If this account is correct, then we should expect to see separate neural signatures of structure learning and associative learning that are systematically related to behavioral performance. However, there is currently little direct neural evidence for structure learning (Collins et al., 2014; Tervo et al., 2016; Madarasz et al., 2016). In this study, we seek to address this gap using human fMRI and an associative learning paradigm adapted from Gershman (2017). On each block, subjects were trained on cue-context-outcome combinations that were consistent with a particular causal interpretation. Subjects were then asked to make predictions about novel cues and contexts without feedback, revealing the degree to which their beliefs conformed to a specific causal structure. We found that the structure learning model developed by Gershman (2017) accounted for the subjects’ predictive judgments, which led us to hypothesize a neural implementation of its computational components.

We found trial-by-trial signals tracking structure learning and associative learning in distinct neural systems. A whole-brain analysis revealed a univariate signature of Bayesian updating of the probability distribution over causal structures in parietal and prefrontal regions, while updating of associative weights recruited a more posterior network of regions. Among these areas, the angular gyrus and the inferior frontal gyrus exhibited structure learning signals that were predictive of subsequent generalization on test trials. Additionally, some of these areas were also implicated in the representation of the full distribution over causal structures. Our results provide new insight into the neural mechanisms of causal structure learning and how they constrain the acquisition of associations.

## Materials and Methods

### Subjects

Twenty-seven healthy subjects were enrolled in the fMRI portion of the study. Although we did not perform power analysis to estimate the sample size, it is consistent with the size of the pilot group of subjects that showed a robust behavioral effect (Figure 4, grey circles), as well as with sample sizes generally employed in the field (Desmond and Glover, 2002). Prior to data analysis, seven subjects were excluded due to technical issues, insufficient data, or excessive head motion. The remaining 20 subjects were used in the analysis (10 female, 10 male; 19-27 years of age; mean age 20 ± 2; all right handed with normal or corrected-to-normal vision). Additionally, 10 different subjects were recruited for a behavioral pilot version of the study that was conducted prior to the fMRI portion. All subjects received informed consent and the study was approved by the Harvard University Institutional Review Board. All subjects were paid for their participation.

### Experimental Design and Materials

We adapted the task used in Gershman (2017) to a within-subjects design. Subjects were told that they would play the role of a health inspector trying to determine the cause of illness in different restaurants around the city. The experiment consisted of nine blocks. Each block consisted of 20 training trials followed by four test trials. On each training trial, subjects were shown a given cue (the food) in a given context (the restaurant) and asked to predict whether that cue-context combination would cause sickness. After making a prediction, they were informed whether their prediction was correct (Figure 1A). On a given block, the assignment of stimuli to outcomes was deterministic, such that the same cue-context pair always led to the same outcome. Even though the computational model could support stochastic and dynamically evolving stimulus-outcome contingencies, our goal was to provide a minimalist design that can assess the main predictions of the theory. There were four distinct training cue-context pairs (two foods × two restaurants) on each block, such that two of the pairs always caused sickness while the other two never caused sickness. Each cue-context pair was shown five times in each block for a total of 20 randomly shuffled training trials.

**Figure 1.**
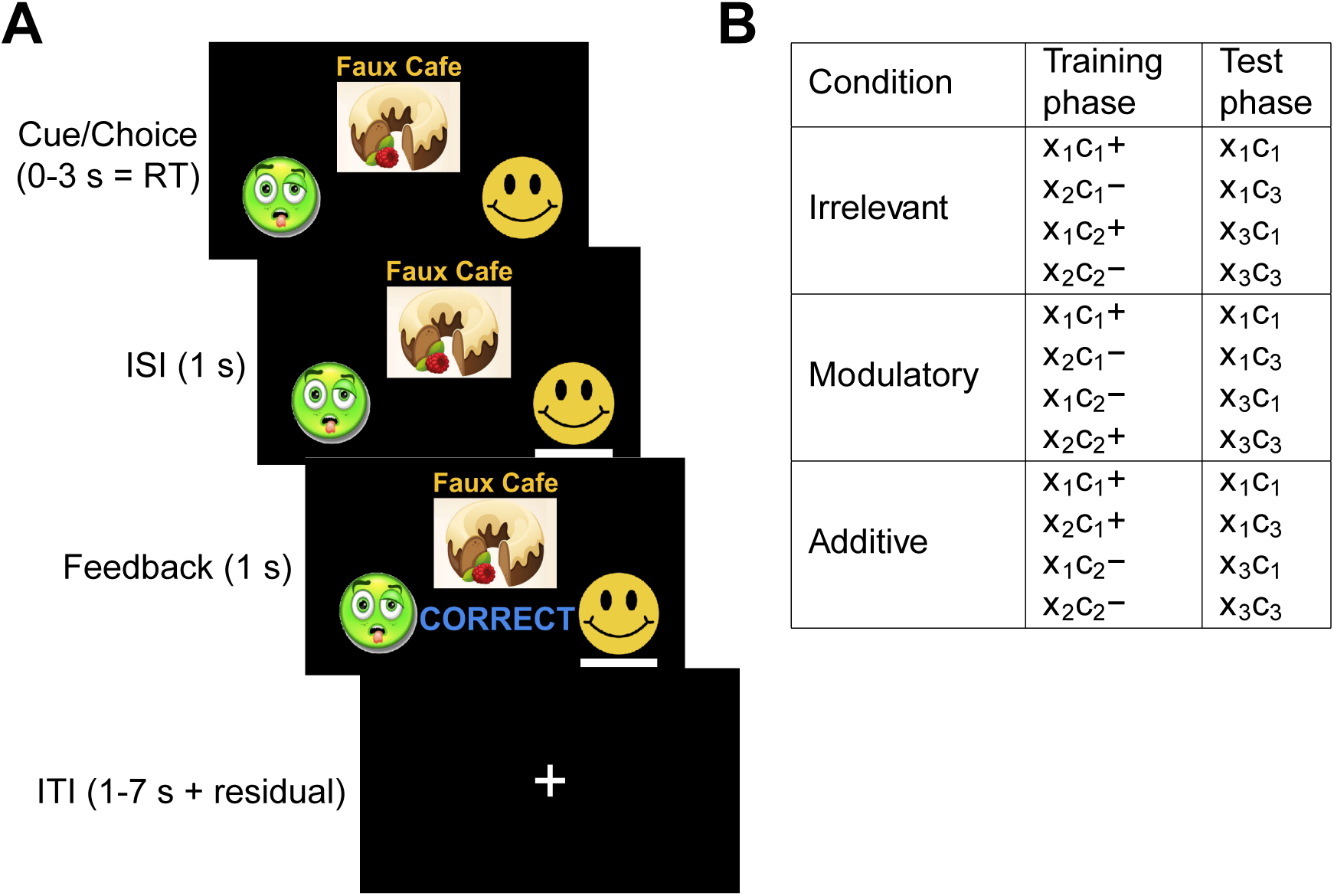
Experimental Design. (A) Timeline of events during a training trial. Subjects are shown a cue (food) and context (restaurant) and are asked to predict whether the food will make a customer sick. They then see a line under the chosen option, and feedback indicating a “Correct” or “Incorrect” response. ISI: interstimulus interval; ITI: intertrial interval. (B) Stimulus-outcome contingencies in each condition. Cues denoted by (*x*_1_, *x*_2_, *x*_3_) and contexts denoted by (*c*_1_,*c*_2_,*c*_3_). Outcome presentation denoted by “+” and no outcome denoted by “−”.

Crucially, the stimulus-outcome contingencies in each block were designed to promote a particular causal interpretation of the environment (Figure 1B, Figure 2). On *irrelevant context* blocks, one cue caused sickness in both contexts, while the other cue never caused sickness, thus rendering the contextual stimulus irrelevant for making correct predictions. On *modulatory context* blocks, the cue-outcome contingency was reversed across contexts, such that the same cue caused sickness in one context but not the other, and vice versa for the other cue. On these blocks, context thus acted like an “occasion setter”, determining the sign of the cue-outcome association. Finally, on *additive context* blocks, both cues caused sickness in one context but neither cue caused sickness in the other context, thus favoring an interpretation of context acting as a punctate cue which sums together with other cues to determine the outcome. There were no explicit instructions or other signals that indicated the different block conditions other than the stimulus-outcome contingencies. While other interpretations of these contingencies are also possible (e.g., that *additive context* sequences also match a causal structure in which cues are irrelevant), we sought to construct the simplest stimulus-outcome relationships that distinguish between the three causal structures outlined in the Introduction. These causal structures correspond to alternative hypotheses about the role of context in associative learning that have been put forward in the animal learning literature (Balsam and Tomie, 1985). We based our experimental design on the fact that a parsimonious model with these structures alone can capture a wide array of behavioral phenomena (Gershman, 2017) and that the chosen stimuli-outcome contingencies establish a clear behavioral pattern that we can build upon to explore the neural correlates of structure learning. The inherent asymmetry between cues and contexts is justified by the *a priori* notion that foods (cues) are spatially and temporally confined focal stimuli, while restaurants (contexts) often comprise spatially and temporally diffuse bundles of background stimuli.

**Figure 2.**
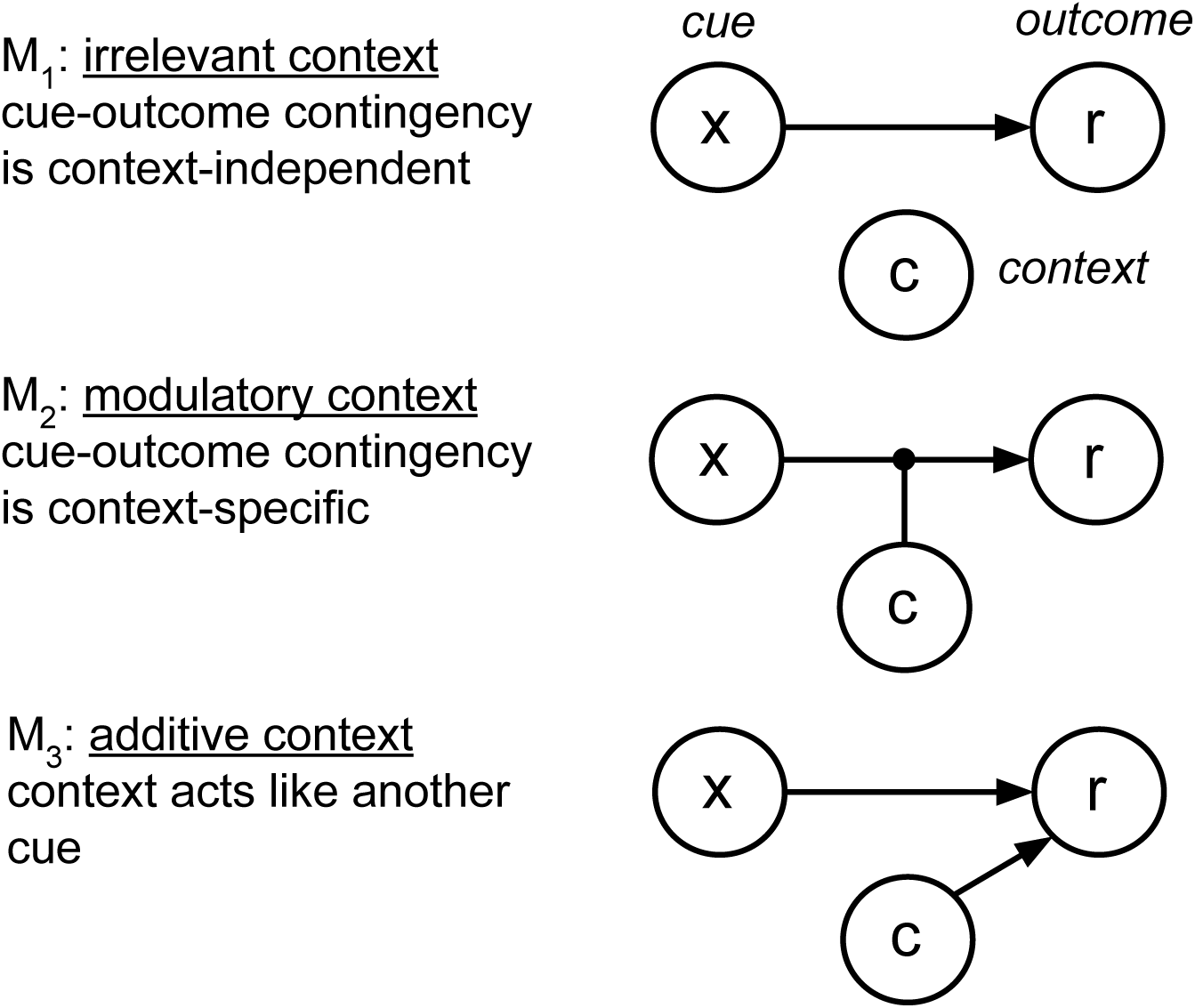
Hypothesis Space of Causal Structures. Each causal structure is depicted as a network where the nodes represent variables and the edges represent causal connections. In *M*_2_, the context modulates the causal relationship between the cue and the outcome. Adapted from Gershman (2017).

Behavior was evaluated on four test trials during which subjects were similarly asked to make predictions, however this time without receiving feedback. Subjects were presented with one novel cue and one novel context, resulting in four (old cue vs. new cue) × (old context vs. new context) randomly shuffled test combinations (Figure 1B). The old cue and the old context were always chosen such that their combination caused sickness during training. Importantly, different causal structures predict different patterns of generalization on the remaining three trials which contain a new cue and/or a new context. If context is deemed to be irrelevant, the old cue should always predict sickness, even when presented in a new context. If a modulatory role of context is preferred, then no inferences can be made about any of the three pairs that include a new cue or a new context. Finally, if context is interpreted as acting like another cue, then both the old cue and the new cue should predict sickness in the old context but not in the new context.

Each block was assigned to one of the three conditions (irrelevant, modulatory, or additive) and each condition appeared three times for each subject, for a total of nine blocks. The block order was randomized in groups of three, such that the first three blocks covered all three conditions in a random order, and so did the next three blocks and the final three blocks. We used nine sets of foods and restaurants corresponding to different cuisines (Chinese, Japanese, Indian, Mexican, Greek, French, Italian, fast food and brunch). Each set consisted of three clipart food images (cues) and three restaurant names (contexts). For each subject, blocks were randomly matched with cuisines, such that subjects had to learn and generalize for a new set of stimuli on each block. The assignment of cuisines was independent of the block condition. The valence of the stimuli was also randomized across subjects, such that the same cue-context pair could predict sickness for some subjects but not others.

### Experimental protocol

Prior to the experiment, the investigator read the task instructions aloud and subjects completed a single demonstration block of the task on a laptop outside the scanner. Subjects completed nine blocks of the task in the scanner, with one block per scanner run. Each block had a duration of 200 seconds during which 100 volumes were acquired (TR = 2 s). At the start of each block, a fixation cross was shown for 10 seconds and the corresponding 5 volumes were subsequently discarded. This was followed by the training phase, which lasted 144 seconds. The event sequence within an example training trial is shown in Figure 1. At trial onset, subjects were shown a food and restaurant pair and instructed to make a prediction. Subjects reported their responses by pressing the left or the right button on a response box. After trial onset, subjects were given 3 seconds to make a response. A response was immediately followed by a 1-second inter-stimulus interval (ISI) during which their response was highlighted. The residual difference between 3 seconds and their reaction time was added to the subsequent inter-trial interval (ITI). The ISI was followed by a 1-second feedback period during which they were informed whether their choice was correct. If subjects failed to respond within 3 seconds of trial onset, no response was recorded and at feedback they were informed that they had timed out. During the ITIs, a fixation cross was shown. The trial order and the jittered ITIs for the training phase were generated using the `optseq2` program (Greve, 2002) with ITIs between 1 and 12 seconds. The training phase was followed by a 4 second message informing the subjects they are about to enter the test phase. The test phase lasted 36 seconds. Test trials had a similar structure as training trials, with the difference that subjects were given 6 seconds to respond instead of 3 and there was no ISI nor feedback period. The ITIs after the first 3 test trials were 2, 4, and 6 seconds, randomly shuffled. The last training trial was followed by a 6-second fixation cross. The stimulus sequences and ITIs were pre-generated for all subjects. The task was implemented using the `PsychoPy2` package (Peirce, 2007). The subjects in the behavioral pilot version of the study performed an identical version of the experiment, except that it was conducted on a laptop.

### Computational modeling

We implemented the model presented in Gershman (2017). The key idea is that learners track the joint posterior over associative weights (**w**) and causal structures (*M*), computed using Bayes’ rule:

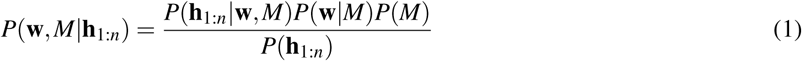

where **h**_1:*n*_ = (**x**_1:*n*_, **r**_1:*n*_, **c**_1:*n*_) denotes the training history for trials 1 to *n* (cue-context-outcome combinations). The likelihood *P*(**h**_1:*n*_|**w**,*M*) encodes how well structure *M* predicts the training history, the prior *P*(**w**|*M*) specifies a preference for weights close to 0, and the prior over structures *P*(*M*) was taken to be uniform. The zero-weight prior assumes a belief that generally most foods do not cause sickness. The structure prior assumes that all structures are equally probable *a priori*.

#### Generative model

Our model is based on the following assumptions about the dynamics that govern associations between stimuli and outcomes in the world. The training history is represented as **h**_1:*n*_ = (**x**_1:*n*_, **r**_1:*n*_, **c**_1:*n*_) for trials 1 to *n*, consisting of the following variables:

- **x**_*n*_ ∈ ℝ^*D*^: the set of *D* cues observed at time *n*, where *x_nd_* = 1 indicates that cue *d* is present and *x_nd_* = 0 to indicate that it is absent. Thus each cue can be regarded as a “one-hot” *D*-dimensional vector and **x**_*n*_ can be viewed as the sum of all cues present on trial *n*. In our simulations, we use *D* = 3 and we only have a single cue (the food) present on each trial.
- *C_n_* ∈ {1,…,*K*}: the context, which can take on one of *K* discrete values. While contexts could in principle be represented as vectors as well, we restrict the model to one context per trial for simplicity. In our simulations, we take *K* = 3.
- *r_n_* ∈ ℝ: the outcome. In our simulations, we use *r_n_* = 1 for “sick” and *r_n_* = 0 for “not sick”.

We consider three specific structures relating the above variables. All the structures have in common that the outcome is assumed to be drawn from a Gaussian with variance 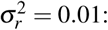

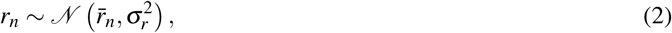

where we have left the dependence on **c**_*n*_ and **x**_*n*_ implicit. The structures differ in how the mean *r̅_n_* is computed.

- **Irrelevant context** (*M*_1_):

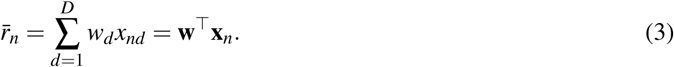

where *d* indexes the set of *D* cues. Under this structure, context *c_n_* plays no role in determining the expected outcome *r̅_n_* on trial *n*. Instead, a single set of weights **w** dictates the associative strength between each cue and the outcome, such that the expected outcome on a given trial is the sum of the associative weights of all present cues. The idea that context is irrelevant for stimulus-outcome associations is consistent with number of behavioral studies (Bouton and King, 1983; Lovibond et al., 1984; Kaye et al., 1987; Bouton and Peck, 1989). For example, if the associative weights are **w** = [1 0.5 0], then cue 1 would predict an average outcome of *r̅_n_* = [1 0.5 0]^┬^[1 0 0] = 1, cue 2 would predict an average outcome of *r̅_n_* = [1 0.5 0]^┬^[0 10]= 0.5, while cue 3 would predict no outcome, on average, since *r̅_n_* = [1 0.5 0]^┬^[0 0 1] = 0. Each cue would make the same prediction regardless of the context *c_n_*, since the weights are the same across contexts.
- **Modulatory context** (*M*_2_):

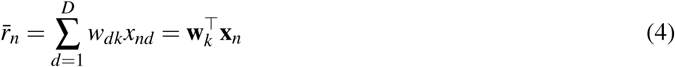

when *c_n_* = *k*. Under this structure, each context *c_n_* = *k* specifies its own weight vector **w**_*k*_. Thus the same cue can make completely different predictions in different contexts. The view that context modulates stimulus-outcome associations is also supported by previous behavioral findings (Swartzentruber and Bouton, 1986; Grahame et al., 1990; Bouton and Bolles, 1993; Swartzentruber, 1995). For example, context 1 may induce associative weights **w**_1_ = [1 0.5 0], while context 2 may induce associative weights **w**_2_ = [0 0.5 1]. Thus in context 2, the predictions of cue 1 and cue 3 would be the opposite of those in context 1.
- **Additive context** (*M*_3_):

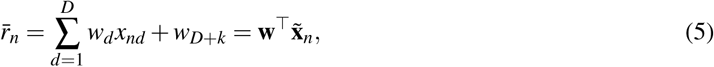

again for *c_n_* = *k*. The augmented stimulus **x̃**_*n*_ is defined as: **x̃**_*n*_ = [**x**_*n*_, **c̃**_*n*_], where *c̃_nk_* = 1 if *c_n_* = *k*, and 0 otherwise. Under this structure, we assume a one-hot vector **c̃**_*n*_ that encodes the context in the same way as the cue. The augmented stimulus **x̃**_*n*_ ∈ ℝ^*D*+*K*^ and the weight vector **w** thus contain entries for both cues and contexts, whose associative strength sums to predict the expected outcome. Previous work also suggests that context sometimes acts like another cue (Balaz et al., 1981; Grau and Rescorla, 1984). For example, imagine that there are *D* = 3 cues and *K* = 3 contexts with associative weights **w** =[10.50-101]. Consider the prediction of cue 1 (**x**_*n*_ = [10 0]) across the three contexts. In context 1, the context vector is **c̃**_*n*_ = [10 0], and hence the augmented stimulus vector is **x̃**_*n*_ = [100 10 0]. In this case, the expected outcome is *r̅_n_* = **w**^┬^**x̃**_*n*_ = 1 × 1 + (−1) × 1 = 0. Even though cue 1 predicts an average outcome of 1 on its own, context 1 predicts an average outcome of −1, and the two cancel each other out to predict no outcome, on average. In context 2, we have **c̃**_*n*_ = [010], **x̃**_*n*_ =[1 000 1 0], and *r̅_n_* = 1 × 1 + 0 × 1 = 1. Context 2 has no associative strength and thus cue 1 predicts an average outcome of 1. In context 3, **c̃**_*n*_ = [0 0 1], **x̃**_*n*_ = [1 0000 1], and *r̅_n_* = 1 × 1 + 1 × 1 = 2. Both cue 1 and context 3 predict an average outcome of 1 separately, so together, they predict double that amount.

We assume each weight is drawn independently from a zero-mean Gaussian prior with variance 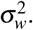 This prior variance was a free parameter that we fit using data from the behavioral pilot version of the study. Each weight can change slowly over time according to a Gaussian random walk with variance *τ*^2^ = 0.001.

In summary, each causal structure corresponds to an internal model of the world in which the relationship between cues, contexts and outcomes can be described by a distinct linear-Gaussian dynamical system (LDS). While the LDS assumptions might seem excessive given the deterministic nature of the task, they have been widely used in the classical conditioning studies (Dayan and Kakade, 2000; Kakade and Dayan, 2002; Kruschke, 2008; Gershman, 2015) to provide a parsimonious account for various learning phenomena. Here we employ them for the purposes of tractability and in order to remain consistent with the causal learning model that Gershman (2017) used to explain the seemingly contradictory roles of context reported in the animal learning literature. The three causal structures correspond to different sub-models that have been advanced in various subforms in the literature, none of which has been able to capture the broad range of results on its own. By recasting each sub-model as an alternative causal structure within the same generative framework, the full model is able to account for a wide range of behavioral findings.

#### Probabilistic inference

Assuming this generative model, a rational agent can use Bayesian inference to invert the model and use its training history **h**_1:*n*_ to learn the underlying causal structure *M* and its associative weights **w** (Eq. 1). To achieve this, first we can compute the posterior over the weights for a given model *M* using Bayes’ rule:

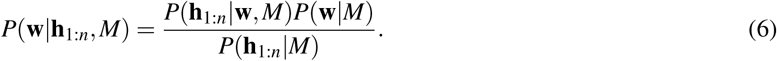

For *M*_1_, the posterior at time *n* is:

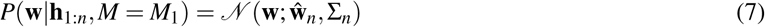

with parameters updated recursively as follows:

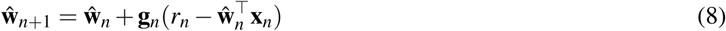

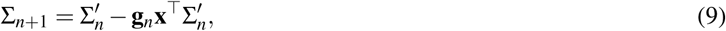

where Σ′_*n*_ = Σ_*n*_ + τ^2^**I**. These update equations are known as *Kalman filtering*, an important algorithm in engineering and signal processing that has recently been applied to animal learning (Dayan and Kakade, 2000; Kruschke, 2008; Gershman, 2015). The initial estimates are given by the parameters of the prior: 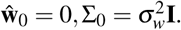 The Kalman gain **g**_*n*_ (a vector of learning rates) is given by:

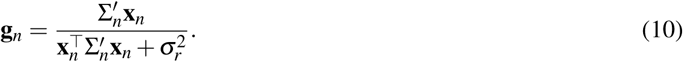

The same equations apply to *M*_2_, but the mean and covariance are context-specific: 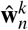 and 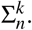 Accordingly, the Kalman gain is modified as follows:

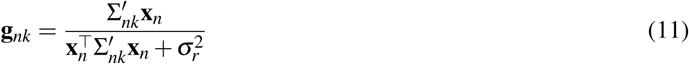

if *c_n_* = *k*, and a vector of zeros otherwise. For *M*_3_, the same equations as *M*_1_ apply, but to the augmented stimulus **x̃**_*n*_.

To make predictions about future outcomes, we need to compute the posterior predictive expectation, which is also available in closed form:

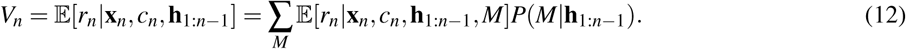

The first term in Eq. 12 is the posterior predictive expectation conditional on model *M*:

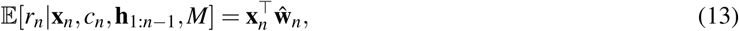

where again the variables are modified depending on what model is being considered. The second term in Eq. 12 is the posterior probability of model *M*, which can be updated according to Bayes’ rule:

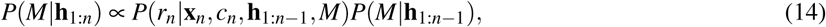

where the likelihood is given by:

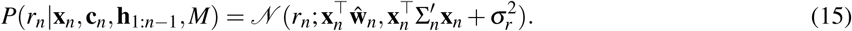

To make predictions for the predictive learning experiment, we mapped the posterior predictive expectation onto choice probability (outcome vs. no outcome) by a logistic sigmoid transformation:

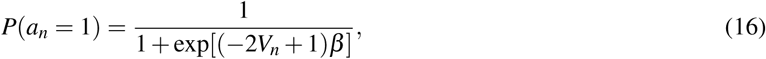

where *a_n_* = 1 indicates a prediction that the outcome will occur, and *a_n_* = 0 indicates a prediction that the outcome will not occur. The free parameter *β* corresponds to the inverse softmax temperature and was fit based on data from the behavioral pilot portion of the study.

In summary, we use standard Kalman filtering to infer the parameters of the LDS corresponding to each causal structure. This yields a distribution over associative weights **w** for each causal structure *M* (Eq. 6), which we can use in turn to compute a distribution over all three causal structures (Eq. 14). The joint distribution over weights and causal structures is then used to predict the expected outcome *V_n_* (Eq. 12) and the corresponding decision *a_n_* (Eq. 16). Our model thus makes predictions about computations at two levels of inference: at the level of causal structures (Eq. 14) and at the level of associative weights for each structure (Eq. 6).

### Parameter estimation

The model has two free parameters: the variance 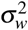 of the Gaussian prior from which the weights are assumed to be drawn, and the inverse temperature *β* used in the logistic transformation from predictive posterior expectation to choice probability. Intuitively, the former corresponds to the level of uncertainty in the initial estimate of the weights, while the latter reflects choice stochasticity. We estimated a single set of parameters based on choice data from the behavioral pilot version of the study using maximum log-likelihood estimation (Figure 4B, grey circles). We preferred this approach over estimating a separate set of parameters for each subject as it tends to avoid overfitting, produces more stable estimates, and has been frequently used in previous studies (Daw et al., 2006; Gershman et al., 2009; Gläscher, 2009; Gläscher et al., 2010). Additionally, since none of these pilot subjects participated in the fMRI portion of the study, this procedure ensured that the parameters used in the final analysis were not overfit to the choices of the scanned subjects. For the purposes of fitting, the model was trained and tested over the same stimulus sequences as the pilot subjects. Each block was simulated independently i.e. the parameters of the model were reset to their initial values prior to the start of training. The likelihood of the subject’s response on a given trial was estimated according to the choice probability given by the model on that trial. Maximum log-likelihood estimation was computed using MATLAB’s fmincon function with 5 random initializations. The bounds on the parameters were 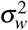 ∈ [0,1] and *β* ∈ [0,10].

**Figure 3.**
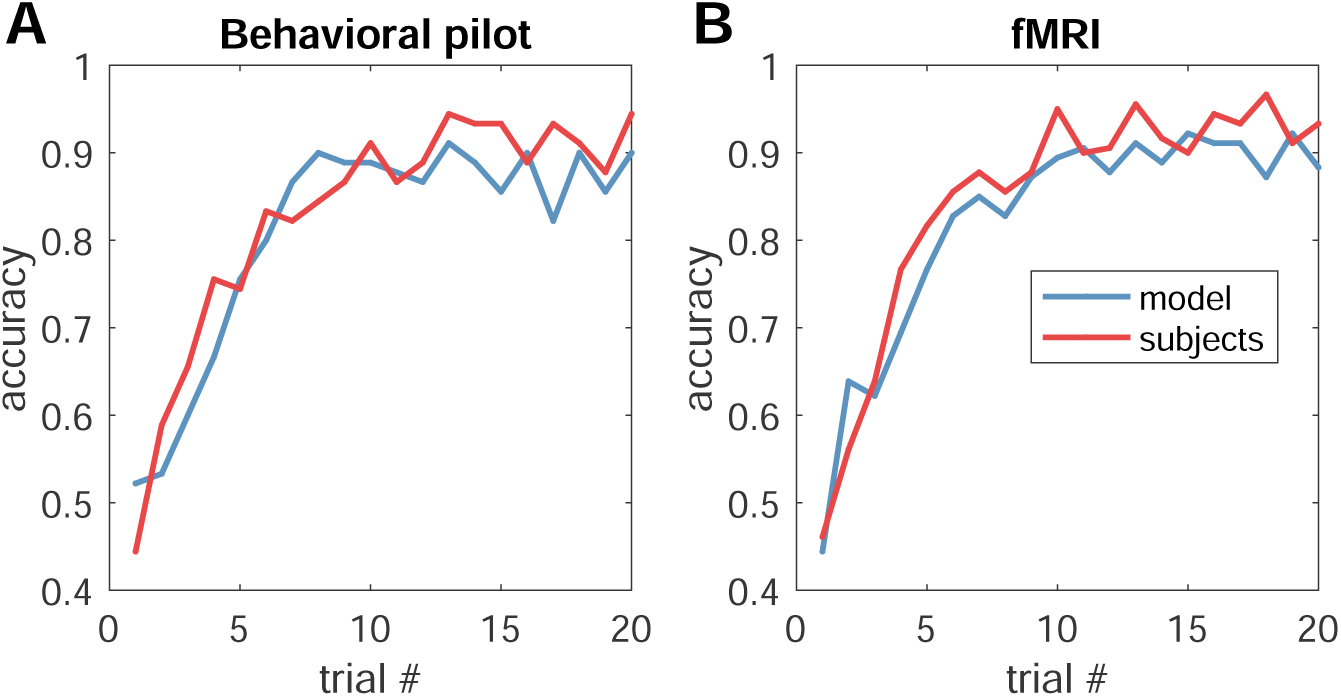
Learning curves during training. Performance during training for (A) behavioral pilot subjects (*N* = 10), and (B) fMRI subjects (*N* = 20), averaged across subjects and blocks. The prior variance of the weights 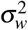 and the logistic inverse temperature *β* were fitted using data only from the pilot version of the study.

**Figure 4.**
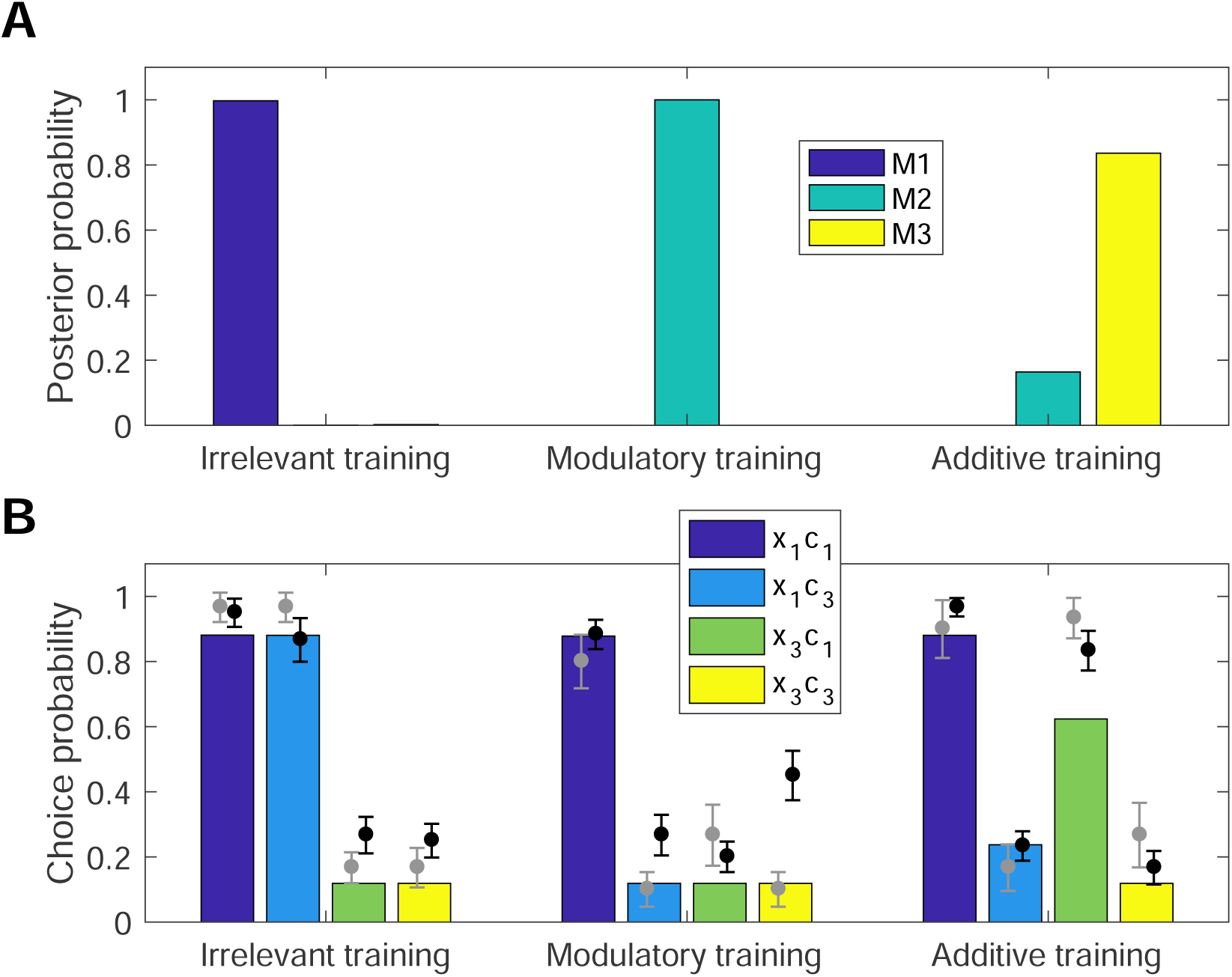
Generalization on the test trials. (A) Posterior probability distribution over causal structures in each condition at the end of training. Each block was simulated independently and the posterior probabilities were averaged across blocks of the same condition. (B) Choice probabilities on the test trials for subjects in the pilot (grey circles) and fMRI (black circles) portions of the study, overlaid over model choice probabilities (colored bars). Each color corresponds to a particular combination of an old (*x*_1_) or new (*x*_3_) cue in an old (*c*_1_) or new (*c*_3_) context. Error bars represent within-subject standard errors of the mean (Cousineau, 2005)

The fitted values were 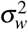 = 0.12 and *β* = 2.01. All other parameters were set to the same values as described in Gershman (2017). We used these parameters to simulate subjects from the fMRI experiment. For behavioral analysis, we trained and tested the model on each block separately and reported the choice probabilities on test trials, averaged across conditions. We used the probability distributions from these simulations to compute the parametric modulators (the Kullback-Leibler divergence) in the GLM (Figure 5) and the representational dissimilarity matrices (Figure 6).

**Figure 5.**
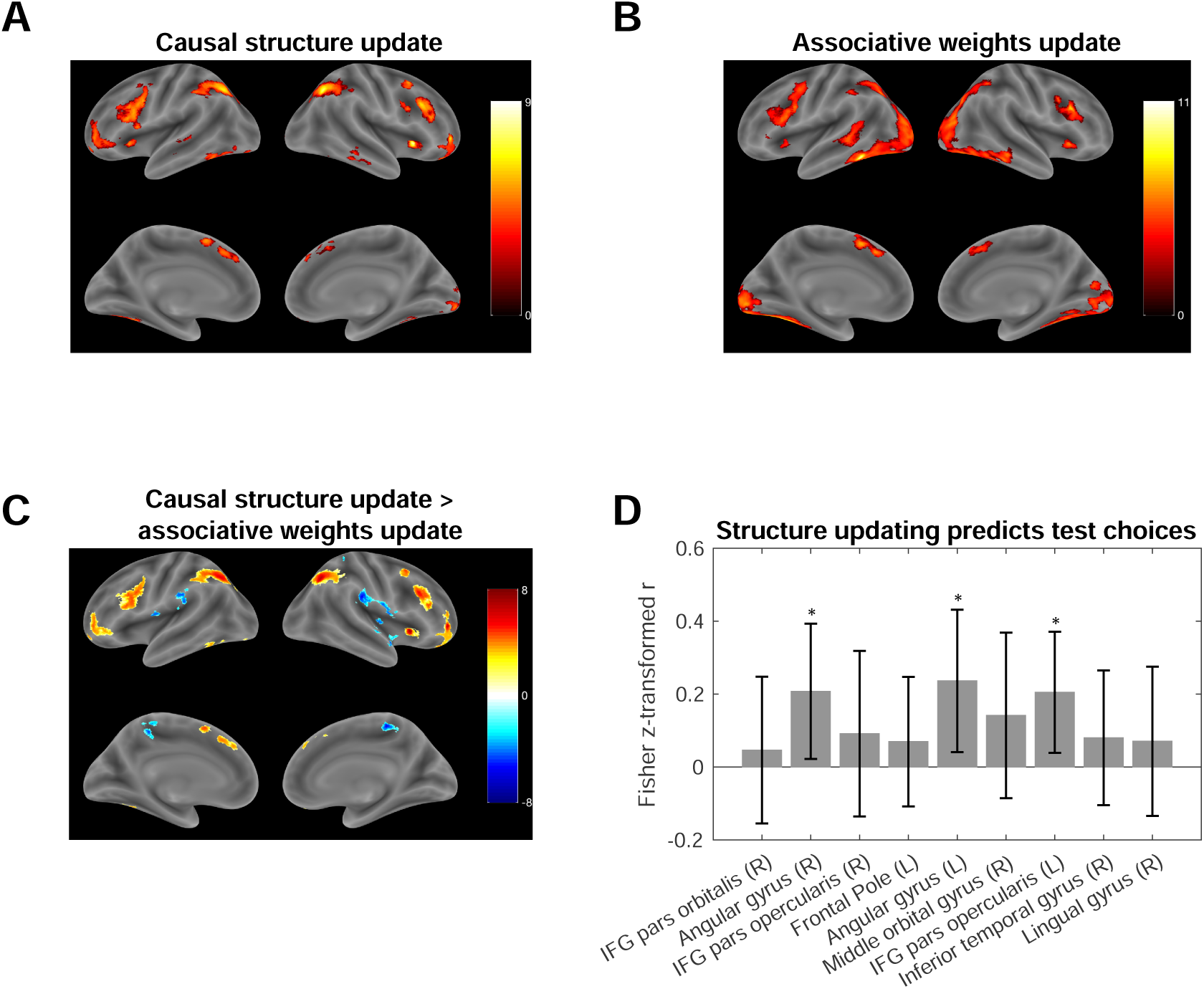
Neural Signature of Structure Learning and Associative Weight Estimation. (A-C) Statistical maps showing regions tracking the KL divergence between the posterior and the prior over causal structures (A) and associative weights (B), and the contrast between the two (C), using a threshold of *p* < 0.001, whole-brain cluster FWE corrected at *α* = 0.05). The color scales represent *t*-values. (D) Bayesian updating of beliefs about causal structure in the angular gyri (MNI: [34,−64,48] and [−30, −62,40]) and in the opercular part of left inferior frontal gyrus (MNI: [−42,2,32]) predict subsequent performance on the test trials. Each bar represents the Fisher *z*-transformed within-subject correlation coefficient between block-specific KL betas and the log likelihoods of the subject’s choices during the test phase (averaged across subjects; error bars: s.e.m.; * = *p* < 0.05).

**Figure 6.**
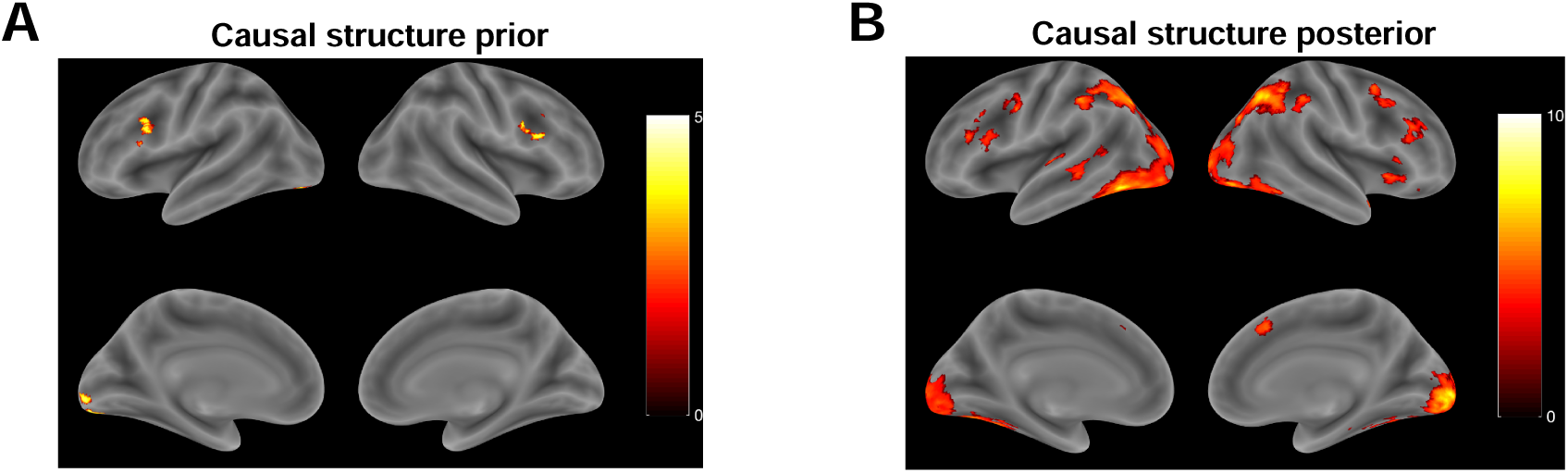
Representational Similarity Analysis. (A, B) Statistical maps showing regions with a high representational similarity match for the distribution over causal structures (*p* < 0.001, whole-brain cluster FWE corrected at *α* = 0.05). The color scales represent *t*-values. (A) Neural activity at trial onset matching the prior. (B) Neural activity at feedback onset matching the posterior.

For model comparison, we similarly estimated parameters for versions of model constrained to use a single causal structure (*M*_1_, *M*_2_, or *M*_3_). We used random effects Bayesian model selection (Rigoux et al., 2014) to compare the three single-structure models and the full model, using −0.5 * BIC as an approximation of the log model evidence, where BIC is the Bayesian information criterion. Models were compared based on their computed exceedance probabilities, with a higher exceedance probability indicating a higher frequency of the given model in the population compared to the other models.

### fMRI data acquisition

Scanning was carried out on a 3T Siemens Magnetom Prisma MRI scanner with the vendor 32-channel head coil (Siemens Healthcare, Erlangen, Germany) at the Harvard University Center for Brain Science Neuroimaging. A T1-weighted high-resolution multi-echo magnetization-prepared rapid-acquisition gradient echo (ME-MPRAGE) anatomical scan (van der Kouwe et al., 2008) of the whole brain was acquired for each subject prior to any functional scanning (176 saggital slices, voxel size = 1.0 × 1.0 × 1.0 mm, TR = 2530 ms, TE = 1.69 − 7.27 ms, TI = 1100 ms, flip angle = 7°, FOV = 256 mm). Functional images were acquired using a T2*-weighted echo-planar imaging (EPI) pulse sequence that employed multiband RF pulses and Simultaneous Multi-Slice (SMS) acquisition (Moeller et al., 2010; Feinberg et al., 2010; Xu et al., 2013). In total, 9 functional runs were collected per subject, with each run corresponding to a single task block (84 interleaved axial-oblique slices per whole brain volume, voxel size = 1.5 × 1.5 × 1.5 mm, TR = 2000 ms, TE = 30 ms, flip angle = 80°, in-plane acceleration (GRAPPA) factor = 2, multi-band acceleration factor = 3, FOV = 204 mm). The initial 5 TRs (10 seconds) were discarded as the scanner stabilized. Functional slices were oriented to a 25 degree tilt towards coronal from AC-PC alignment. The SMS-EPI acquisitions used the CMRR-MB pulse sequence from the University of Minnesota. Four subjects failed to complete all 9 functional runs due to technical reasons and were excluded from the analyses. Three additional subjects were excluded due to excessive motion.

### fMRI preprocessing

Functional images were preprocessed and analyzed using `SPM12` (Wellcome Department of Imaging Neuroscience, London, UK). Each functional scan was realigned to correct for small movements between scans, producing an aligned set of images and a mean image for each subject. The high-resolution T1-weighted ME-MPRAGE images were then co-registered to the mean realigned images and the gray matter was segmented out and normalized to the gray matter of a standard Montreal Neurological Institute (MNI) reference brain. The functional images were then normalized to the MNI template (resampled voxel size 2 mm isotropic), spatially smoothed with a 8 mm full-width at half-maximum (FWHM) Gaussian kernel, high-pass filtered at 1/128 Hz, and corrected for temporal autocorrelations using a first-order autoregressive model.

### Univariate analysis

To analyze the fMRI data, we defined a GLM with four impulse regressors convolved with the canonical hemodynamic response function (HRF): a regressor at trial onset for all training trials (the stimulus regressor); a regressor at feedback onset for all training trials (the feedback regressor); a regressor at feedback onset for training trials on which the subject produced the wrong response (the error regressor); and a regressor at feedback onset for training trials on which the contextual stimulus differs from the contextual stimulus on the previous trial (the context regressor). The context regressor was not included on the first training trial of each block. The feedback regressor had two parametric modulators: the Kullback-Leibler (KL) divergence between the posterior and the prior probability distribution over causal structures; and the KL divergence between the posterior and the prior probability distribution over the associative weights for the causal structure corresponding to the condition of the given block. In particular, the causal structure KL divergence on trial *n* was computed as:

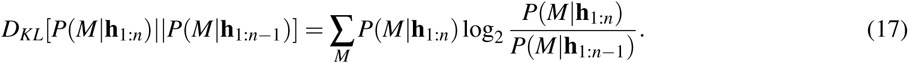

The associative weights KL divergence for trial *n* and causal structure *M*_1_ was computed as:

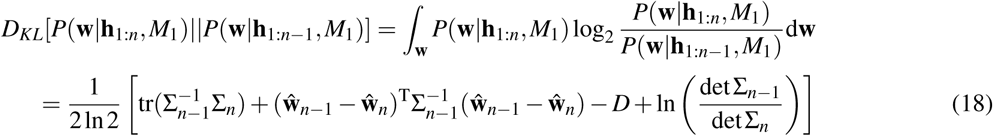

where *D* denotes the number of weights, Σ_*n*_ denotes the posterior covariance on trial *n*, and dividing by ln2 converts the result to bits. Eq. 18 follows from the fact that the weights are normally distributed (Eq. 7). The KL divergence was computed analogously for causal structures *M*_2_ and *M*_3_. The two parametric modulators were not orthogonalized with respect to each other nor with respect to the feedback regressor. Orthogonalizing the two KL divergences would introduce an ordering effect which would unnecessarily complicate the analysis. Furthermore, computing the contrast between the two parametric modulators (Figure 5C) would subtract any common variability in the signal.

For group-level analyses, we report *t*-contrasts with single voxels thresholded at *p* < 0.001 and whole-brain cluster family-wise error (FWE) correction applied at significance level *α* = 0.05. Anatomical regions were labeled using the Automated Anatomical Labeling (AAL2) atlas (Tzourio-Mazoyer et al., 2002; Rolls et al., 2015), the SPM Anatomy Toolbox (Eickhoff et al., 2005), and the CMA Harvard-Oxford atlas (Desikan et al., 2006). All voxel coordinates are reported in Montreal Neurological Institute (MNI) space.

### Correlating neural activity with behavior

In order to link neural variability in structure updating with behavior, we identified the peak voxel (i.e. voxel with the highest *t*-statistic) in each activation cluster from the contrast corresponding to the KL divergence for causal structures (Figure 5A and Table 2). For each peak voxel, we defined a region of interest (the peak ROI) as a 27-voxel sphere (radius = 1.814 voxels) centered on that voxel and intersected with the corresponding activation cluster (i.e., only voxels that belong both to the sphere and to the cluster were included in the ROI). These ROIs represent the distinct brain areas that are most strongly correlated with causal structure updating. For each block and for each subject, we then extracted the structure KL betas of the voxels in each ROI. Since the betas are computed for each block separately as an intermediate step in fitting the GLM, we did not have to perform any additional analyses in order to obtain them.

For a given block, the structure KL beta of a particular voxel represents the degree to which BOLD activity in that voxel tracks the causal structure KL divergence across all 20 training trials of the block. Even for voxels that are on average highly correlated with the KL divergence (such as the peak voxels), there is block-to-block variability in the strength of that relationship. If the activity of a given voxel represents updates about causal structures, a lower beta on a particular block would suggest that the underlying computations deviated from Bayesian updating as predicted by the model. Conversely, a high beta would indicate that the computations were consistent with the structure learning account. Thus one would expect that performance on the test phase following a given block would be similar to the predictions of the Bayesian model in the latter case but not in the former. We tested this idea by correlating the average beta across all voxels in the peak ROI for a given training block with the average log likelihood of the subject’s choices on the test trials following that block. This yielded a Pearson’s correlation coefficient (*n* = 9 blocks per subject) for each subject. The resulting correlation coefficients were Fisher *z*-transformed and entered into a one-sample *t*-test against 0. For each peak ROI, we report the average Fisher *z*-transformed *r* and the result from the *t*-test (Figure 5D).

Primary sensory, visual, and motor regions such as V1 and the cerebellum were *a priori* excluded from this analysis. Negative activation clusters were also excluded. We chose spherical ROIs of this size in order to remain consistent with our searchlight analysis and with previous studies (Chan et al., 2016). Similarity between the model predictions and the subject’s behavior was computed as the mean log likelihood of their choices across test trials on which they produced a response, 〈log*P*(*c_n_*)〉, where *c_n_* = 1 indicates that the subject predicted the outcome would occur on trial *n*, *c_n_* = 0 indicates that they predicted the outcome would not occur, and the probability *P*(*c_n_*) is calculated as in Eq. 16.

### Representational similarity analysis

We used representational similarity analysis (RSA) to identify candidate brain regions that might encode the full probability distribution over causal structures in their multivariate activity patterns (Kriegeskorte et al., 2008; Xue et al., 2010; Chan et al., 2016). On a given trial, we expected Bayesian updating to occur when the outcome of the subject’s prediction is presented at feedback onset (i.e. whether they were correct or incorrect). We therefore expected to find representations of the prior distribution at trial onset and representations of the posterior distribution at feedback onset (which in turn corresponds to the prior distribution on the following trial). We describe how we performed RSA for the prior; the analysis for the posterior proceeded in a similar fashion.

In order to identify regions with a high representational similarity match for the prior, we used an unbiased whole-brain “searchlight” approach. For each voxel of the entire volume, we defined a 27-voxel spherical ROI (radius = 1.814 voxels) centered on that voxel, excluding voxels outside the brain. For each subject and each ROI, we computed a 180 × 180 representational dissimilarity matrix **R** (the neural RDM) such that the entry in row *i* and column *j* is the cosine distance between the neural activity patterns on training trial *i* and training trial *j*:

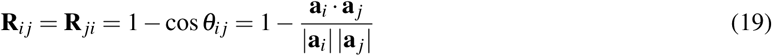

where **θ**_*ij*_ is the angle between the 27-dimensional vectors **a**_*i*_ and **a**_*j*_ which represent the instantaneous neural activity patterns at trial onset on training trials *i* and *j*, respectively, in the given ROI for the given subject. Neural activations entered into the RSA were obtained using a GLM with distinct impulse regressors convolved with the HRF at trial onset and feedback onset on each trial (test trials had regressors at trial onset only). The neural activity of a given voxel was thus simply its beta coefficient of the regressor for the corresponding trial and event. Since the matrix is symmetric and **R**_*ii*_ = 0, we only considered entries above the diagonal (i.e. *i* < *j*). The cosine distance is equal to 1 minus the normalized correlation (i.e. the cosine of the angle between the two vectors), which has been preferred over other similarity measures as it better conforms to intuitions about similarity both for neural activity and for probability distributions (Chan et al., 2016).

Similarly, we computed an RDM (the model RDM) such that the entry in row *i* and column *j* is the cosine distance between the priors on training trial *i* and training trial *j*, as computed by model simulations using the stimulus sequences experienced by the subject on the corresponding blocks.

If neural activity in the given ROI encodes the prior, then the neural RDM should resemble the model RDM: trials on which the prior is similar should have similar neural representations (i.e. smaller cosine distances), while trials on which the prior is dissimilar should have dissimilar neural representations (i.e. larger cosine distances). This intuition can be formalized using Spearman’s rank correlation coefficient between the model RDM and the neural RDM (*n* = 180 × 179/2 = 16110 unique pairs of trials in each RDM). A high coefficient implies that pairs of trials with similar priors tend show similar neural patterns while pairs of trials with dissimilar priors tend to show dissimilar neural patterns. Spearman’s rank correlation is a preferred method for comparing RDMs over other correlation measures as it does not assume a linear relationship between the RDMs (Kriegeskorte et al., 2008). Thus for each voxel and each subject, we obtained a single Spearman’s *ρ* that reflects the representational similarity match between the prior and the ROI around that voxel.

In order to aggregate these results across subjects, for each voxel we Fisher *z*-transformed the resulting Spearman’s *ρ* from all 20 subjects and performed a *t*-test against 0. This yielded a group-level *t*-map, where the *t*-value of each voxel indicates whether the representational similarity match for that voxel is significant across subjects. We thresholded single voxels at *p* < 0.001 and corrected for multiple comparisons using whole-brain cluster FWE correction at significance level *α* = 0.05. We report the surviving clusters and the *t*-values of the corresponding voxels (Figure 6A).

We used an identical procedure to obtain a *t*-map indicating which brain regions how a high representational similarity match with the posterior (Figure 6B). The only difference was that we computed the neural RDMs using activity patterns at feedback onset (rather than at trial onset) and we computed the model RDMs using the posterior (instead of the prior).

Finally, since both the prior and the posterior tend to be similar on trials that are temporally close to each other, as well as on trials from the same block, we computed two control RDMs: a “time RDM” in which the distance between trials *i* and *j* is |*t_i_* − *t_j_*|, where *t_i_* is the difference between the onset of trial *i* and the start of its corresponding block; and a “block RDM” in which the distance between trials *i* and *j* is 0 if they belong to the same block, and 1 otherwise. Each Spearman’s *ρ*’s was then computed as a partial rank correlation coefficient between the neural RDM and the model RDM, controlling for the time RDM and the block RDM. This rules out the possibility that our RSA results reflect within-block temporal autocorrelations that are unrelated to the prior or the posterior.

## Results

### Causal structure learning accounts for behavioral performance

The behavioral results replicated the findings of Gershman (2017) using a within-subject design. Subjects from both the pilot and the fMRI portions of the study learned the correct stimulus-outcome associations relatively quickly, with average performance plateauing around the middle of training (Figure 3). Average accuracy during the second half of training was 91.2 ± 2.5% (*t*_9_ = 16.8, *p* < 10^−7^, one-sample *t*-test against 50%) for the pilot subjects, and 92.7 ± 1.7% (*t*_19_ = 25.0, *p* < 10^−15^, one-sample *t*-test against 50%) for the scanned subjects, well above chance.

Importantly, both groups exhibited distinct patterns of generalization on the test trials across the different conditions, consistent with the results of Gershman (2017) (Figure 4B). Without taking the computational model into account, these generalization patterns already suggest that subjects learned something beyond simple stimulus-response mappings On blocks during which context was irrelevant (Figure 4B, irrelevant training), subjects tended to predict that the old cue *x*_1_, which caused sickness in both *c*_1_ and *c*_2_, would also cause sickness in the new context *c*_3_ (circle for *x*_1_*c*_3_), even though they had never experienced *c*_3_ before. The new cue *x*_3_, on the other hand, was judged to be much less predictive of sickness in either context (*t*_38_ = 9.51,*p* < 10^−10^, paired *t*-test). Conversely, on blocks during which context acted like another cue (Figure 4B, additive training), subjects guessed that both cues would cause sickness in the old context *c*_1_ (circle for *x*_3_*c*_1_), but not in the new context *c*_3_ (*t*_38_ = 11.1, *p* < 10^−12^, paired *t*-test). Generalization in both of these conditions was different from what one would expect if subjects treated each cue-context pair as a unique stimulus independent from the other pairs, which is similar to the generalization pattern on modulatory blocks (Figure 4B, modulatory training). On these blocks, subjects judged that the old cue is predictive of sickness in the old context significantly more compared to the remaining cue-context pairs (*t*_38_ = 9.01, *p* < 10^−10^, paired *t*-test).

These observations were consistent with the predictions of the computational model. Using parameters fit with data from the behavioral pilot version of the study, the model quantitatively accounted for the generalization pattern on the test trials choices of subjects in the fMRI portion of the study (Figure 4B; *r* = 0.96,*p* < 10^−6^). As expected, the stimulus-outcome contingencies induced the model to infer a different causal structure in each of the three conditions (Figure 4A), leading to the distinct response patterns on the simulated test trials. For comparison, we also ran versions of the model using a single causal structure. Theories corresponding to each of these sub-models have been put forward in the literature as explanations of the role of context during learning, however neither of them has been able to provide a comprehensive account of the behavioral findings on its own. Accordingly, model performance was markedly worse when the hypothesis space was restricted to a single causal structure: the correlation coefficients were *r* = 0.61 for the irrelevant context structure (*M*_1_; *p* = 0.04), *r* = 0.73 for the modulatory context structure (*M*_2_; *p* < 0.01), and *r* = 0.91 for the additive context structure (*M*_3_; *p* < 0.0001). Bayesian model comparison confirmed the superiority of the full structure learning model over the restricted variants (Table 1).

**Table 1.** Model comparison. Each model is represented by its corresponding hypothesis space of causal structures (Figure 2). The free parameters 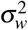 and *β* were fit based on choice data from the pilot version of the study (Figure 4B, grey circles). Log likelihood and Pearson’s r were computed based on choice data from the fMRI version of the study. Bayesian information criterion (BIC) and protected exceedance probability (PXP) were computed on the pilot data. Log lik, log likelihood.

| Hypotheses | $\sigma_w^2$ | $\beta$ | BIC | PXP | Log lik | Pearson's r |
| --- | --- | --- | --- | --- | --- | --- |
| $M_1, M_2, M_3$ | 0.1249 | 2.0064 | 1670 | 0.3855 | -1711 | $r = 0.96, p < 10^{-6}$ |
| $M_1$ | 0.0161 | 1.2323 | 2631 | 0.2048 | -2687 | $r = 0.61, p = 0.036$ |
| $M_2$ | 0.9997 | 1.7433 | 1828 | 0.2049 | -1895 | $r = 0.73, p = 0.008$ |
| $M_3$ | 0.0130 | 1.7508 | 2184 | 0.2048 | -2180 | $r = 0.91, p = 0.00004$ |

### Separate brain regions support structure learning and associative learning

We sought to identify brain regions in which the blood oxygenation level dependent (BOLD) signal tracks beliefs about the underlying causal structure. In order to condense these multivariate distributions into scalars, we computed the Kullback-Leibler (KL) divergence between the posterior and the prior distribution over causal structures on each training trial, which measures the degree to which structural beliefs were revised after observing the outcome. Specifically, we analyzed the fMRI data using a general linear model (GLM) which included the KL divergence as a parametric modulator at feedback onset. We reasoned that activity in regions involved in learning causal structure would correlate with the degree of belief revision.

A high KL divergence often implies that, before receiving feedback on the current trial, the agent had inferred the wrong causal structure. Accordingly, we found that the KL divergence was relatively higher for trials on which subjects produced an incorrect response (two-sample *t*-test, *t*_3598_ = 12.2, *p* < 10^−30^). To eliminate any confounds specifically related to receiving negative feedback about incorrect responding, we included an error regressor at feedback onset indicating trials on which subjects made an incorrect prediction. In addition, since the structures are sensitive to contextual stimuli by design, the KL divergence tended to be higher when context changed from the previous trial (two-sample *t*-test, *t*_3418_ = 6.53,*p* < 10^−10^). In order to account for potential effects of contextual tracking, we thus included a regressor at feedback onset on trials on which the contextual stimulus differs from the previous trial. Finally, since we were interested in regions that correlate with learning on the level of causal structures rather than their associative weights, we included the KL divergence between the posterior and the prior distribution over associative weights for the causal structure corresponding to the condition of the given block (e.g., only the weights for the modulatory structure were analyzed on modulatory blocks). These weights encode the strength of causal relationships between cues and outcomes (and additionally context, in the case of the additive structure). Including this KL divergence as an additional parametric modulator at feedback onset would capture any variability in the signal related to weight updating and allow us to isolate it from the signal related to structure updating.

We computed three group-level contrasts: the KL divergence for causal structures, the KL divergence for associative weights, and the difference between the two. Activation clusters in all contrasts were identified by thresholding single voxels at *p* < 0.001 and applying whole-brain cluster FWE correction at significance level *α* = 0.05. The first contrast revealed several regions that were associated with updating of the posterior over causal structures (Figure 5A, and Table 2). We observed bilateral parietal clusters with peak activations in the angular gyri. We also found bilateral clusters in inferior frontal gyrus (IFG) spanning the opercular and the triangular parts of IFG. Bilateral activations were also found in rostrolateral PFC, extending into the medial and anterior orbital gyri in the right hemisphere and into the anterior insula and the orbital part of IFG in the left hemisphere. There was a cluster around the orbital part of IFG in the right hemisphere as well. Significant activations were also found in bilateral cerebellum, bilateral supplementary motor area (SMA), and in left inferior and middle temporal gyrus.

**Table 2.** Brain regions in which the BOLD signal tracks the KL divergence between the posterior and the prior over causal structures (corresponding to Figure 5A). The voxel with the maximum *t*-statistic from each cluster is also reported. Single voxels were thresholded at *p* < 0.001 and whole-brain cluster FWE correction was applied at significance level *α* = 0.05. IFG, inferior frontal gyrus. MNI, Montreal Neurological Institute. BA, Brodmann area.

| Sign | Brain region | BA | Extent | *t*-value | MNI coord. |
| --- | --- | --- | --- | --- | --- |
| Positive | IFG pars orbitalis (R) | 47 | 263 | 9.474 | 32 22 0 |
|  | Angular gyrus (R) | 39 | 1539 | 8.687 | 34 -64 48 |
|  | IFG pars opercularis (R) | 44 | 1411 | 7.483 | 48 20 32 |
|  | Rostrolateral PFC (L) | 10 | 1305 | 7.406 | -40 56 6 |
|  | Angular gyrus (L) | 39 | 1500 | 7.343 | -30 -62 40 |
|  | Rostrolateral PFC (R) | 10 | 789 | 7.296 | 36 54 0 |
|  | IFG pars opercularis (L) | 44 | 1560 | 6.999 | -42 2 32 |
|  | Supplementary motor area (L) | 6 | 868 | 6.286 | -6 8 52 |
|  | Cerebelum (R) |  | 660 | 6.217 | 18 -70 -24 |
|  | Cerebelum (L) |  | 1378 | 5.913 | -6 -76 -28 |
|  | Cerebelum (L) |  | 207 | 5.646 | -32 -68 -52 |
|  | Inferior temporal gyrus (R) | 20 | 159 | 5.533 | 62 -24 -22 |
|  | Lingual gyrus (R) | 19 | 225 | 5.364 | 12 -96 -8 |
|  | Middle temporal gyrus (L) | 21 | 187 | 4.367 | -58 -36 -4 |
| Negative | Lateral ventricle (L) |  | 425 | -8.748 | -26 -36 12 |
|  | Insula (R) | 13 | 1935 | -7.211 | 46 -8 6 |
|  | Insula (L) | 13 | 300 | -6.907 | -42 -14 0 |
|  | Precuneus (R) | 7 | 443 | -6.513 | 10 -40 52 |
|  | Supramarginal gyrus (L) | 40 | 455 | -6.020 | -66 -32 36 |
|  | Precuneus (L) | 7 | 368 | -6.019 | -12 -42 46 |

The second contrast revealed a more posterior network of brain regions that was correlated with the KL divergence for associative weights (Figure 5B, and Table 3). We found a large cluster spanning visual cortex bilaterally and extending into inferior parietal and inferior temporal cortex. We observed bilateral activations in IFG that were similar to those observed in the causal structures KL contrast. There were also activations in the orbital part of bilateral IFG, left middle temporal gyrus, and bilateral SMA. We also found subcortical activations in right thalamus and bilateral ventral striatum.

**Table 3.** Brain regions in which the BOLD signal tracks the KL divergence between the posterior and the prior distribution over associative weights (corresponding to Figure 5B). Notation and procedures as in Table 2.

| Sign | Brain region | BA | Extent | *t*-value | MNI coord. |
| --- | --- | --- | --- | --- | --- |
| Positive | Inferior temporal, parietal,<br>and occipital gyri (L) |  | 12042 | 11.091 | -50 -52 -16 |
|  | IFG pars triangularis (R) | 45 | 645 | 7.935 | 36 20 26 |
|  | IFG pars opercularis (L) | 44 | 1663 | 7.929 | -42 6 26 |
|  | Supplementary motor area (L) | 6 | 1187 | 7.452 | -8 10 50 |
|  | IFG pars orbitalis | 47 | 181 | 7.334 | 30 26 0 |
|  | Middle temporal gyrus (L) | 21 | 542 | 6.088 | -52 -38 -2 |
|  | Thalamus and Striatum (R) |  | 421 | 6.009 | 12 -14 2 |
|  | IFG pars orbitalis (L) | 47 | 528 | 5.269 | -40 20 -4 |
| Negative | Lateral ventricle (L) |  | 314 | -7.449 | 24 -46 16 |
|  | Postcentral gyrus (R) | 5 | 390 | -6.962 | 26 -46 62 |
|  | Lateral ventricle (R) |  | 466 | -6.786 | -20 -46 12 |
|  | Precuneus (L) | 7 | 572 | -6.261 | -26 -48 66 |
|  | Postcentral gyrus (R) | 1 | 1293 | -5.853 | 60 -12 34 |

In order to isolate regions that are specifically related to causal structure updating and not associative weight updating, we computed the contrast between these two regressors (Figure 5C, and Table 4). Positive activations in this contrast correspond to areas where the signal is more strongly correlated with the causal structures KL divergence than with the associative weights KL divergence, while negative activations indicate the opposite relationship. This contrast highlighted essentially the same areas as the single regressor contrast with the causal structures KL divergence (Figure 5A, and Table 2), implying that the signal in these areas cannot be explained by associative learning alone and suggesting a dissociable network of regions that might support structure learning. It is worth noting that the regions which correlated more strongly with associative weight updating were in bilateral insula, bilateral precuneus, and left supramarginal gyrus. Inspecting the single regressor contrasts revealed that these regions were either not significantly correlated or were negatively correlated with both regressors. This suggests that their negative activation here is due to a more negative correlation with the causal structures KL divergence, rather than to a positive correlation with the associative weights KL divergence or a common representation of both types of updating.

**Table 4.** Brain regions in which the BOLD signal is better explained by the causal structures KL divergence than by the associative weights KL divergence (corresponding to Figure 5C). Notation and procedures as in Table 2.

| Sign | Brain region | BA | Extent | *t*-value | MNI coord. |
| --- | --- | --- | --- | --- | --- |
| Positive | IFG pars orbitalis (R) | 47 | 263 | 9.474 | 32 22 0 |
|  | Angular gyrus (R) | 39 | 1539 | 8.687 | 34 -64 48 |
|  | Rostrolateral PFC (R) | 10 | 746 | 7.948 | 36 54 0 |
|  | Rostrolateral PFC (L) | 10 | 881 | 7.045 | -40 56 6 |
|  | Angular gyrus (L) | 39 | 1500 | 7.343 | -30 -62 40 |
|  | Middle frontal gyrus (R) | 9 | 1096 | 6.666 | 40 32 22 |
|  | IFG pars opercularis (L) | 44 | 1049 | 6.159 | -38 14 26 |
|  | Supplementary motor area (L) | 6 | 543 | 5.758 | -6 8 52 |
|  | Cerebelum (R) |  | 304 | 5.498 | 18 -70 -24 |
|  | Cerebelum (L) |  | 394 | 5.417 | -8 -78 -30 |
|  | Cerebelum (L) |  | 165 | 5.194 | -32 -68 -52 |
|  | Fusiform gyrus (L) | 37 | 206 | 4.705 | -42 -52 -14 |
| Negative | Lateral ventricle (L) |  | 327 | -8.327 | -26 -36 12 |
|  | Insula (R) | 13 | 937 | -6.712 | 46 -8 6 |
|  | Insula (L) | 13 | 169 | -6.290 | -42 -14 0 |
|  | Supramarginal gyrus (L) | 40 | 298 | -5.876 | -66 -32 36 |
|  | Precuneus (R) | 7 | 182 | -5.758 | 10 -40 52 |
|  | Temporal pole (R) | 38 | 200 | -5.419 | 50 4 -14 |
|  | Precuneus (L) | 7 | 192 | -5.381 | -12 -42 46 |

### Neural correlates of causal structure updating during training predict subsequent test performance

We hypothesized that if a neural signal correlates with updating beliefs about causal structure during training, it should also be associated with variability in behavior during the test trials. For each block and for each subject, we computed the average parameter estimate of the causal structure KL regressors across voxels in a 27-voxel spherical region of interest (ROI) centered on the peak voxel from each activation cluster (i.e. the peak voxels from Table 2; see Materials and Methods). Intuitively, these averaged betas (the KL betas) correspond to how strongly the voxels from a given ROI correlate with the causal structure KL divergence during the training trials of a given block. If a particular region mediates structure learning, then variability in its KL beta should predict variability in test phase behavior across blocks. Accordingly, for each subject and each ROI, we correlated the KL beta across blocks with the log likelihood of the subject’s responses on the test trials given the model predictions. The test log likelihood indicates how closely the subject’s responses conform to the model predictions. For example, if on a particular block subjects are updating their structural beliefs, then their test phase performance should more closely resemble the predictions of the structure learning model, which would be reflected in a high test log likelihood. Thus for a given ROI, a positive correlation across subjects would indicate that the extent to which that region tracks Bayesian updating during training predicts the extent to which test performance is consistent with causal structure learning. Importantly, since the KL betas were estimated on the training trials preceding the test phase, this analysis would yield an unbiased assessment of the neural predictions.

For each subject and each ROI, we thus obtained a within-subject Pearson correlation coefficient across blocks, indicating how well the degree to which that ROI encodes structure updating predicts test choices for that subject. For each ROI, we performed a one-sample *t*-test against 0 with the Fisher *z*-transformed Pearson’s *r* from all subjects (Figure 5D). We found a significant positive relationship in right angular gyrus (*t*_19_ = 2.34, *p* = 0.03; mean *z* = 0.21 ± 0.19; peak MNI coordinates [34 −64 48]), left angular gyrus (*t*_19_ = 2.53,*p* = 0.02, mean *z* = 0.24± 0.20; peak MNI coordinates [−30 −62 40]), and the opercular part of left IFG (*t*_19_ = 2.6, *p* = 0.02, mean *z* = 0.20 ± 0.17; peak MNI coordinates [-42 2 32]). The other regions were not significant (*p*’s > 0.1). These results link neural variability in correlates of Bayesian updating to individual variability in test phase choices, providing a more stringent criterion for identifying the substrates underlying causal structure learning. We performed a similar analysis with the associative weights KL regressor and found no significant relationship (all *p*’s > 0.1).

### Multivariate representations of the posterior and prior distribution over causal structures

If the brain performs Bayesian inference over causal structures, as our data suggest, then we should be able to identify regions that contain representations of the full probability distribution over causal structures before and after the update. We thus performed a whole-brain “searchlight” representational similarity analysis (Kriegeskorte et al., 2008; Xue et al., 2010; Chan et al., 2016) using 27-voxel spherical searchlights. For each subject, we centered the spherical ROI on each voxel of the whole-brain volume and computed a representational dissimilarity matrix (RDM) using the cosine distance between neural activity patterns at trial onset for all pairs of trials (see Materials and Methods). Intuitively, this RDM reflects which pairs of trials look similar and which pairs of trials look different according to the neural representations in the area around the given voxel. We then used Spearman’s rank correlation to compare this neural RDM with a model RDM based on the prior distribution over causal structures. If a given ROI encodes the prior, then pairs of trials on which the prior is similar would also show similar neural representations, while pairs of trials on which the prior is different would show differing neural representations. This corresponds to a positive rank correlation between the model and the neural RDMs.

For each subject and each voxel, we thus obtained a Spearman’s rank correlation coefficient, reflecting the similarity between activity patterns around that voxel and the prior over causal structures (the representational similarity match). For each voxel, we then performed a one-sample *t*-test against 0 with the Fisher *z*-transformed Spearman’s *ρ* from all subjects. The resulting *t*-values from all voxels were used to construct a whole-brain *t*-map (Figure 6A and Table 5). This revealed bilateral activations in PFC near the triangular and opercular parts of IFG, as well as a region in left inferior temporal cortex, suggesting that these areas contain multivariate representations of the prior.

**Table 5.** Brain regions with a high representational similarity match between neural patterns at trial onset and the prior over causal structures (corresponding to Figure 6A). The voxel with the maximum *t*-statistic from each cluster is also reported. Single voxels were thresholded at *p* < 0.001 and whole-brain cluster FWE correction was applied at significance level *α* = 0.05. Notation as in Table 2.

| Sign | Brain region | BA | Extent | $t$-value | MNI coord. |
| --- | --- | --- | --- | --- | --- |
| Positive | Fusiform gyrus (L) | 37 | 279 | 5.687 | -22 -84 -4 |
|  | IFG pars opercularis (L) | 44 | 315 | 5.435 | -36 12 24 |
|  | IFG pars triangularis (R) | 45 | 279 | 5.276 | 48 18 26 |

We performed the same analysis using neural activations at feedback onset and the posterior over causal structures (Figure 6B and Table 6). This revealed a broader set of regions, including a large cluster in visual cortex that extended bilaterally into inferior temporal cortex and the angular gyri, overlapping with the regions found in the univariate analyses. We also found bilateral activations near the triangular and opercular parts of IFG, as well as a small cluster around the orbital part of right IFG. Taken together, these results suggest that the brain maintains representations of the full probability distribution over causal structures, which it updates in accordance with Bayesian inference on a trial-by-trial basis as new evidence is accumulated.

**Table 6.**
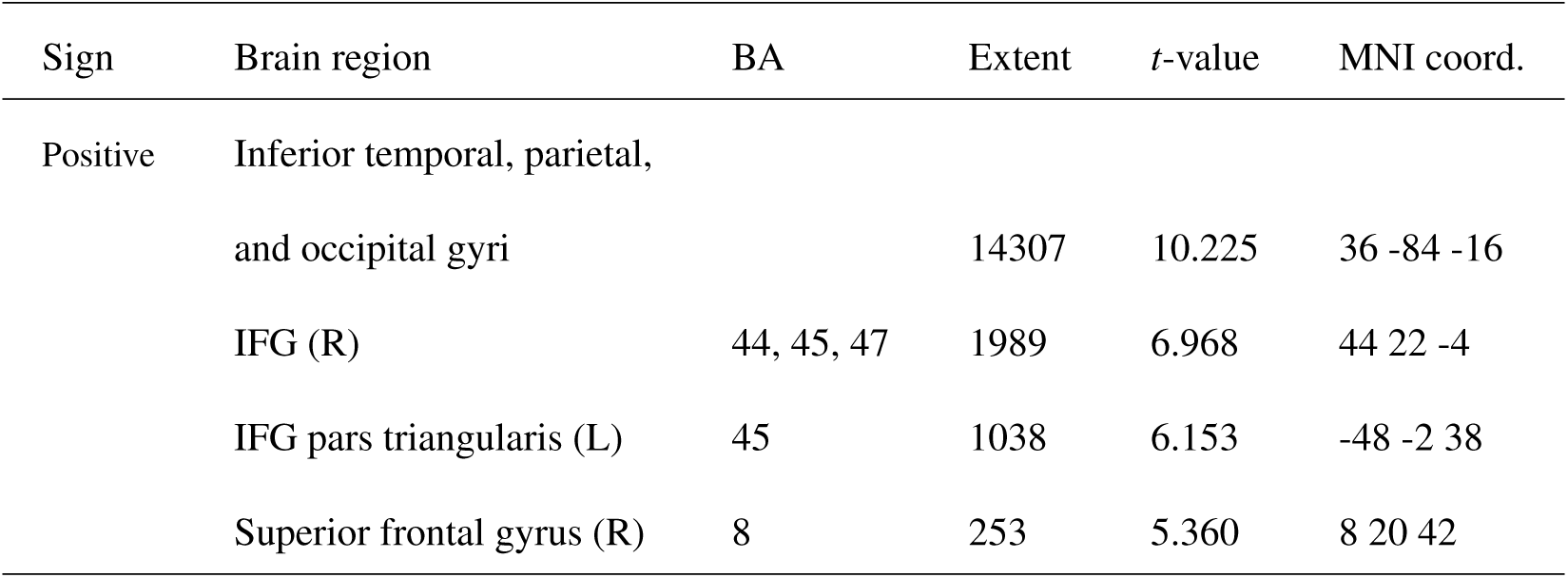
Brain regions with a high representational similarity match between neural patterns at feedback onset and the posterior over causal structures (corresponding to Figure 6B). Notation and procedures as in Table 5.

| Sign | Brain region | BA | Extent | *t*-value | MNI coord. |
| --- | --- | --- | --- | --- | --- |
| Positive | Inferior temporal, parietal,<br>and occipital gyri |  | 14307 | 10.225 | 36 -84 -16 |
|  | IFG (R) | 44, 45, 47 | 1989 | 6.968 | 44 22 -4 |
|  | IFG pars triangularis (L) | 45 | 1038 | 6.153 | -48 -2 38 |
|  | Superior frontal gyrus (R) | 8 | 253 | 5.360 | 8 20 42 |

## Discussion

Behavioral evidence suggests that humans and animals infer both the structure and the strength of causal relationships (Griffiths and Tenenbaum, 2005; Körding et al., 2007; Meder et al., 2014; Gershman, 2017). Using functional brain imaging in humans, the current study provides neural evidence for a causal structure learning account of the role of contextual influences on the formation of stimulus-outcome associations. The neural data support the existence of separate learning mechanisms operating over structural and associative representations, thus reifying the computationally hypothesized division of labor. A univariate analysis identified areas that were sensitive to belief updates about causal structure, including regions of prefrontal, orbitofrontal and parietal cortex. Among these regions, activation in both angular gyri (AG) and in left inferior frontal gyrus (IFG) predicted variability in test phase behavior across blocks. Furthermore, these regions were found to be distinct from the correlates of associative learning, which included inferior temporal and visual areas. In addition, a representational similarity analysis revealed brain areas that appear to represent the full probability distribution over causal structures. Regions representing the posterior overlapped with regions involved in updating the posterior, including bilateral AG and IFG. Representation of the prior at trial onset also appeared in bilateral IFG.

These results build on previous work that has implicated some of these areas in different forms of learning and decision making. We found activations related to weight updating in regions commonly linked to reinforcement learning, such as the ventral striatum, an area that has been associated with the prediction error signal in temporal difference reinforcement learning (O’Doherty et al., 2003; Gläscher et al., 2010). In our model, associative learning also involves the computation of an error term (the difference between the observed and the expected outcome; see Materials and Methods), which is used to revise the posterior distribution over weights.

Our finding that activity in inferior parietal cortex correlates with both kinds of learning could be related to previous reports of elevated AG activity for relational associations (Schwartz et al., 2011; Boylan et al., 2017), evidence accumulation (d’Acremont et al., 2013), and insights (Aziz-Zadeh et al., 2009). The AG has also been implicated in reorienting attentional selection based on task history (Taylor et al., 2011), responding to salient new information (Singh-Curry and Husain, 2009), and bottom-up attentional shifts towards relevant stimuli that were previously ignored (Downar et al., 2000; Ciaramelli et al., 2008). Gläscher et al. (2010) showed that AG encodes the state prediction error in model-based reinforcement learning, reflecting the discrepancy between the current model and the observed state transitions. Another notable study by O’Reilly et al. (2013) found that AG encodes the discrepancy between the prediction of a prospective model that combines prior experience with a dynamic feed-forward simulation and the prediction of a retrospective model that is based on task history alone, which they operationalized as the KL divergence between the two outcome probability distributions. The same study found that AG also tracks the KL divergence between the posterior and the prior outcome probability distributions predicted by retrospective model. AG has also been implicated in schema memory (Gilboa and Marlatte, 2017), particularly in encoding new associations into a schema (van Buuren et al., 2014). Seghier (2013) has attempted to unify the diverse roles of AG by proposing that the region acts as an integrator of bottom-up multimodal input and top-down predictions from frontal areas, with the resulting output then fed back to adjust the higher-level predictions. Our finding that AG activates for both structure updating and weight updating (albeit in anatomically distinct loci), and predicts subsequent choice behavior, fits with the idea that AG acts as a cross-modal hub that integrates prior knowledge with incoming information.

One candidate region where such top-down predictions might originate is lateral prefrontal cortex, an area with strong functional connectivity with the inferior parietal lobule (IPL, which includes AG) (Vincent et al., 2008; Boorman et al., 2009). Previous studies on cognitive control (Koechlin et al., 2003; Koechlin and Summerfield, 2007; Badre and D’Esposito, 2007) have proposed the existence of a functional gradient in lateral PFC, with more anterior regions encoding representations of progressively higher levels of abstraction, culminating in rlPFC. Donoso et al. (2014) found evidence that rlPFC performs inference over multiple counterfactual strategies by tracking their reliability, while the triangular part of IFG is responsible for switching to one of those strategies if the current one is deemed unreliable. If causal structures are likened to alternative strategies, then these results may relate to our finding that rlPFC tracks structure updating, while IFG tracks both structure and weight updating. These updates can be interpreted as discrepancies indicating that the current causal structure and/or its associative weights are unreliable, thus warranting a revision of the weights or a switch to an alternative causal structure.

Our results also resonate with follow-up work on hierarchical reinforcement learning (Badre et al., 2010; Frank and Badre, 2012), which extends the notion of a functional hierarchy in lateral PFC to the acquisition of abstract latent rules that guide stimulus-outcome associations. Frank and Badre (2012) proposed a Bayesian mixture of experts (MoE) model which arbitrates between hypotheses about stimulus features that are relevant for predicting the outcome, assigning greater weight to experts that make better predictions. Most pertinent to our study is a model developed by Collins and Frank (2016) that groups different mappings between stimulus features and responses into distinct task sets. Different contextual stimuli can elicit different task sets, akin to the occasion-setting function of modulatory context in our study.

Despite the apparent similarities, there are several notable distinctions between our theoretical framework and those previous modeling efforts. The term “structure” is used with a different meaning in those studies, either referring to the hierarchical structure of the hypothesis space (Frank and Badre, 2012), or to the mapping between contextual stimuli and task sets (Collins and Frank, 2016). While the model of Collins and Frank (2016) can support an infinite number of task sets, it does not model summation across cues, and therefore cannot represent the additive causal structure from our study. Some form of summation is widely believed to be important for capturing classic animal learning phenomena such as blocking, overshadowing, and overexpectation (Rescorla and Wagner, 1972; Soto et al., 2014).

Our results point to potential avenues for future research that could further interrogate the functional role of the regions we identified. For example, if AG supports causal structure updating, as our data suggest, then temporarily disrupting it during the first half of training (e.g., using transcranial magnetic stimulation) should result in the same response pattern on the test trials across all conditions, since subjects would not be able to update their structural beliefs and to infer different causal structures in the different task conditions. On the other hand, disrupting AG during the second half of training should have little impact on test responses, as learning has usually plateaued by then. Conversely, if the areas identified in lateral PFC maintain the full distribution over causal structures, then disrupting them during the first half of training may slow down learning but would not necessarily result in uniform test responding, as subjects would still be able to catch up during the second half of training. However, disrupting those regions during the second half of training could affect the learned distribution over causal structures and result in uniform test response patterns across all conditions.

An important question that remains open is how structure learning might be implemented in biologically realistic neural circuits. Tervo et al. (2016) noted the parallels between the hierarchical architecture of cortical circuits and the hierarchical nature of structure learning, with empirical evidence suggesting that different layers of the hierarchy tend to be associated with separate cortical circuits. If the brain indeed performs Bayesian inference over causal structures, this raises the more fundamental question of how ensembles of neurons could represent and perform operations on probability distributions. Different theories have been put forward, ranging from probabilistic population codes to Monte Carlo sampling (Pouget et al., 2013). Teasing apart the different possible mechanisms would require developing behavioral frameworks that lend themselves to computational modeling and quantitative predictions about the inferred probability distributions (Tervo et al., 2016). We believe this study is an important step in that direction.

In summary, we used a combination of behavioral, neural and computational techniques to tease apart the neural substrates of structure learning and associative learning. Inference over the space of causal structures recruited prefrontal regions that have been previously implicated in higher-order reasoning and state representations (rlPFC and OFC), whereas learning the associative weights activated regions that have been implicated in reward-based learning (ventral striatum). Both forms of learning elicited bilateral activations in IFG and AG, a region involved in integrating multimodal sensory input with top-down predictions. Moreover, activity in the voxels that correlated most strongly with causal structure updating in AG and IFG predicted the similarity between subsequent subject behavior and the predictions of the causal structure learning model. Finally, representational similarity analysis revealed regions that encode the full distribution over causal structures: a multivariate signature of the prior was found in IFG, while the posterior was associated with a broader set of brain areas, including IFG, AG, and inferior temporal cortex. Together, these results provide strong support for the idea that the brain performs probabilistic inference over latent structures in the environment, enabling inductive leaps that go beyond the given observations.

## Acknowledgements

This research was supported by the National Institutes of Health, award number 1R01MH109177. This work involved the use of instrumentation supported by the NIH Shared Instrumentation Grant Program grant number S10OD020039. We acknowledge the University of Minnesota Center for Magnetic Resonance Research for use of the multiband-EPI pulse sequences. We are grateful to Erik Kastman and Katie Insel for helping to design the experiment and collect the data.

